# Recruitment of transcriptional effectors by Cas9 creates cis regulatory elements and demonstrates distance-dependent transcriptional regulation

**DOI:** 10.1101/2022.02.03.478957

**Authors:** Jubran Boulos, Itai Ehrlich, Nili Avidan, Omer Shkedi, Lilac Haimovich-Caspi, Noam Kaplan, Izhak Kehat

**Affiliations:** The Rappaport Institute and the Bruce Rappaport Faculty of Medicine, Technion – Israel Institute of Technology, 1 Efron Street, P.O. Box 9697 Haifa 3109601, Israel

**Keywords:** Enhancer, Cardiac enhancer, Gene expression, CRISPR, Cis regulation

## Abstract

It is essential to regulate the expression of genes, such as those encoding the proteins of the cardiac sarcomere. This regulation is often mediated by *cis* regulatory elements termed enhancers and repressors that recruit transcription factors to gene-distal sites. However, the relationship between transcription factors recruitment to gene-distant sites and the regulation of gene expression is not fully understood. Specifically, it is unclear if such recruitment to any genomic site is sufficient to form an enhancer or repressor at the site, and what is the relationship between the *cis* regulatory element’s position and its ability to control the transcription of distant genes. Using dead Cas9 to recruit either viral or endogenous transcription factor activation domains, we demonstrate that targeting ‘naïve’ genomic sites lacking open chromatin or active enhancer marks is sufficient to alter the chromatin signature of the target site, the distant gene promoter, and significantly induce the distant gene expression, even across chromatin insulating loci. The magnitude of induction is affected by the distance between the activation site and the cognate gene in a non-linear manner. Dead Cas9 mediated recruitment of repression domains behave similarly to activation in that targeting of non-regulatory regions could repress gene expression with a nonlinear distance dependence and across chromatin insulating loci. These findings expand the models of enhancer generation and function by showing that an arbitrary genomic site can become a regulatory element and interact epigenetically and transcriptionally with a distant promoter. They also provide new fundamental insights into the rules governing gene expression.

## Introduction

Cells can control the expression of their genes through complex regulatory networks, which include the target gene and its regulators. *Cis*-acting regulatory elements (CREs) like enhancers and repressors control distant genes’ expression and play a critical role in gene expression (Field and Adelman, 2020; Rosa-Garrido et al., 2018). The understanding of gene regulation has advanced significantly, but many questions remain unanswered, especially regarding the relationship between transcription factor recruitment to gene-distant sites and the regulation of gene expression. Particularly, it is unclear whether such recruitment to any genomic site is sufficient to form a cis-regulatory element at the site and control the expression of distant genes, and how the *cis*-regulatory element’s position affects its ability to do so.

Recently, the CRISPR/Cas9 system (clustered regularly interspaced short palindromic repeats and CRISPR-associated protein 9) has been repurposed to modulate endogenous gene expression. CRISPR activators (CRISPRa), composed of a nuclease-dead mutant of Cas9 (dCas9) tethered to various transcription factor activator domains were used to induce gene expression (Chavez et al., 2016; Cheng et al., 2013; Gilbert et al., 2014, 2013; Kearns et al., 2014; Konermann et al., 2015; Lin et al., 2015; Maeder et al., 2013; Mali et al., 2013; Perez-Pinera et al., 2013; Simeonov et al., 2017; Tanenbaum et al., 2014). Of those, A hybrid dCas9-VP64-p65-Rta tripartite activator (dCas9-VPR) showed a strong, synergistic activation of several genes, including cardiac genes, when targeted to their promoters (Chavez et al., 2015). Two large screens based on VP64 activation domains and pooled tiling gRNA libraries showed enrichment in gRNAs targeting the proximal promoter near the transcription start sites (TSS) (Gilbert et al., 2014; Simeonov et al., 2017) or in gRNAs targeting regulatory elements marked by open chromatin and Histone 3 Lysine 27 acetylation (H3K27ac) (Simeonov et al., 2017). A dCas9 fused to the p300 histone acetyltransferase domain could activate genes when targeted to enhancers, while dCas9-VP64 did not (Hilton et al., 2015). Collectively, these studies showed that CRISPRa can activate genes and identify regulatory elements but implied that the efficiency of these tools is limited to targeting promoters and enhancers.

Similarly, CRISPR inhibitors (CRISPRi) were developed by tethering the krüppel-associated box (KRAB) repression domain to dCas9 (Fulco et al., 2019, 2016; Gao et al., 2014; Klann et al., 2017; Thakore et al., 2015; Xie et al., 2017). Screens that used dCas9-KRAB and tiling pools of gRNA (Fulco et al., 2016; Gilbert et al., 2014) or pools of gRNA directed at DNase hypersensitive sites (Klann et al., 2017; Xie et al., 2017) showed that gRNAs targeting proximal promoters or enhancers were enriched. As with CRISPRa, these studies showed that CRISPRi could repress genes or screen for regulatory elements, but implied that the efficacy of these tools was limited to targeting promoters and enhancers.

CREs such as enhancers and repressors are binding sites for a collection of transcription factors that together modulate the activity of distant genes (Field and Adelman, 2020). We hypothesized that CRISPRa and CRISPRi can be used to simulate the *de novo* creation of a CRE because they allow the recruitment of multiple transcription factor activation or repressor domains to specific sites in the genomic context. We thus set out to identify the requirements and consequences of such *de novo* CRE generation and to determine the effects of the position of the CRE on the transcriptional output of distant genes. To this end we used CRISPRa to activate cardiac genes in fibroblasts, where they are not normally expressed, and CRISPRi to repress them in cardiomyocytes, where they are highly expressed. We systematically targeted dCas9 effectors to multiple sites within a 140 Kbp window in 6 genomic loci and show that recruiting either viral or endogenous transcription factor effector domains using dCas9 to a naïve genomic site is sufficient to alter the chromatin marks of the targeted site, the distant gene promoter, and change the distant gene expression even from distances of up to 70 Kbp away. This transcriptional control can cross neighboring genes and chromatin insulating loci, and act regardless of the existence of open chromatin at the targeted sites. The effects are non-linearly dependent on the distance and are often stronger when targeting sites close to the gene, providing a model of CRE generation and function.

## Results

### Cardiac enhancers are distributed around cardiac specific genes

To analyze the distribution of endogenous regulatory elements relative to genes, we mapped the position of cardiac-specific regulatory elements relative to cardiac-specific genes. We previously used H3K27ac ChIP-seq, ATAC-seq, and RNA-seq in fibroblasts (FIB) and cardiomyocytes (CM) to identify cell-type-specific regulatory elements and genes (Golan-Lagziel et al., 2018). From these data, we used the RNA-seq to concentrate on the transcription start sites (TSS) of 221 highly cardiac-specific genes, defined as genes with more than 1K read counts whose expression is at least 8-fold higher in CM than in FIB, with adjusted p<0.05. We then mapped the density of CM-specific open regulatory elements, defined by 4-fold enrichment of ATAC-seq tag counts with a Poisson enrichment p-value < 0.0001 in CM vs. FIB, around the TSS of these 221 CM specific genes (Fig. 1a-b). We find that cardiac-specific regulatory elements are densely distributed within 10 Kbp of the TSS of cardiac-specific genes, with a non-linear decline in density as the distance from the TSS increases.

**Fig. 1.**
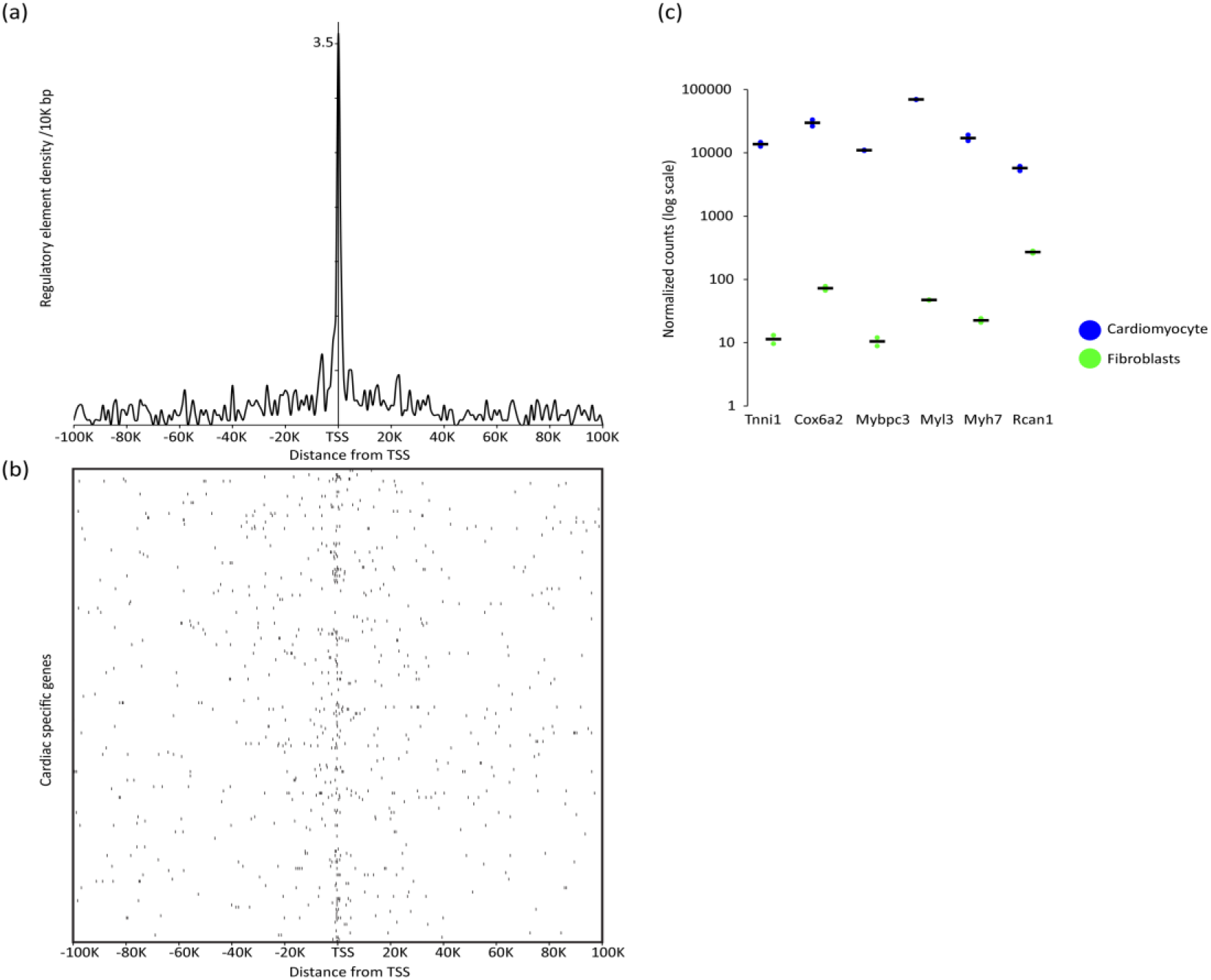
cardiac-specific open regulatory elements are distributed around cardiac specific genes. (**a**) A density histogram of CM-specific open chromatin distribution around the transcription start site (TSS) of 221 cardiac specific genes, showing high density of cardiac specific regulatory elements very close to the TSS with nonlinear decay at greater distances (Kbp). (**b**) Same data as in (a) presented as a map where the 221 loci were centered around the cardiac specific gene TSS and the distribution of CM-specific open chromatin was plotted in a window of ± 100 Kbp. (**c**) Analysis of expression of six selected cardiac specific genes in cardiomyocytes (blue) and fibroblasts (green) using DE-seq2 normalized RNA-seq counts on a logarithmic scale, showing two orders of magnitude higher expression in cardiomyocytes for these genes (n=3 for cardiomyocytes and n=2 for fibroblasts, black bars show the mean expression).

Studies with CRISPRa showed that activation of weakly expressed genes results in higher fold induction (Chavez et al., 2016; Konermann et al., 2015), but that the degree of activation is highly variable among genes (Lin et al., 2015; Mali et al., 2013; Perez-Pinera et al., 2013). Therefore, to map the activation range of dCas9-VPR we looked for genes that are highly expressed in CM, lowly expressed in FIB, that could be robustly activated by targeting dCas9-VPR to their promoter in FIB. From the 221 genes cardiac specific genes we chose six such CM specific genes (*Mybpc3, Myh7, Tnni1, Cox6a2, Myl3*, and *Rcan1*) whose expression is much higher in CM than in FIB (Fig. 1c).

### Recruitment of activation domains upregulates gene expression in a distance dependent manner from multiple genomic loci

To identify the requirements and consequences of de novo CRE creation and to study the effect of the position of the CRE on the transcriptional output, we activated the six CM-specific genes in FIB with dCas9-VPR. Since these genes and the cardiac transcription factors controlling them are very lowly expressed in FIB, these loci provide a background with a minimal regulatory complexity for investigating the consequences of activation domain recruitment. To avoid bias and to systematically cover these loci we chose multiple gRNA target sites for activation in each locus, based solely on distance of the target site from the index gene TSS and on a predicted ability to specifically and efficiently recruit Cas9. Sites were chosen with higher density near the TSS, and with subsequent spacing steps of 5-20 Kbp (Table S1). Both CRISPRa and CRISPRi were previously used with multiplexed gRNAs with no loss of specificity (Konermann et al., 2015; Wang et al., 2019; Zhao et al., 2018). We therefore multiplexed gRNAs for *Mybpc3, Tnni1*, and *Rcan1* loci or for *Myh7, Cox6a2*, and *Myl3*, with a single gRNA for each locus. In both triplexes the targeted loci are each located on different chromosomes. FIBs were transfected with a complex of dCas9-VPR plasmid and site-specific gRNAs triplex. FIB transfected with the same complex but with a non-targeting gRNA served as controls, and qRT-PCR analysis was performed 24 Hrs after transfection to measure gene expression.

We measured the degree of transcriptional activation of *Mybpc3, Myh7, Tnni1, Cox6a2, Myl3*, and *Rcan1* resulting from dCas9-VPR recruitment to multiple sites, spanning 140 Kbp around the TSS, in each of these loci. In total 149 genomic sites were targeted by gRNAs, with 127 (85.2%) of those significantly eliciting a change in target gene expression. These data were used to generate activation maps for each of the six loci, where the fold activation of the index gene above the control is shown as a black bar over the site of the targeting gRNA (Fig. 2a, Fig. S1). In the *Mybpc3, Tnni1, Myl3*, and *Rcan1* loci the activation was strongest from sites close to the TSS (59±32,29±5.5,13.4±4.4, or 6.4±0.57 fold respectively). In the *Cox6a2* locus strong activation was achieved by targeting dCas9-VPR close to the TSS (22.7±1.15 fold) but also at several sites within 30 Kbp down- and up-stream of the TSS (e.g., activation of 16±1.2 and 30.6±3.2 was achieved by targeting sites 20 Kbp upstream and downstream of the TSS respectively). In the *Myh7* locus only low activation was achieved by targeting multiple sites near the TSS, but strong activation (56±10.6 fold) was achieved by targeting a site 20 Kbp downstream, that is closer to the *Myh6* gene promoter. A summary of the degree of activation achieved in all six loci as a function of the distance from the TSS is shown in Fig. 2b.

**Fig. 2.**
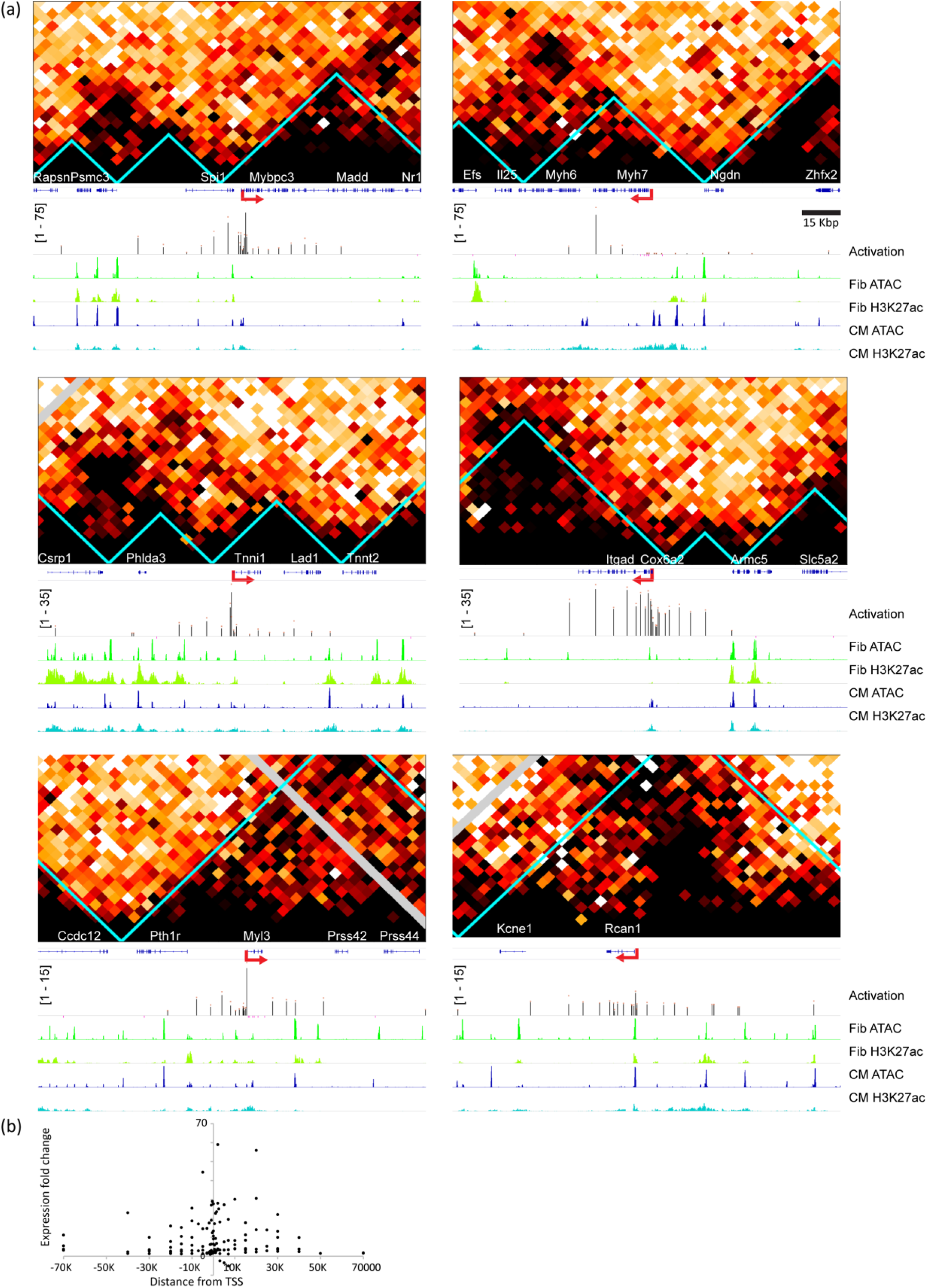
Recruitment of activation domain upregulates gene expression in a distance dependent manner from multiple genomic sites. (**a**) Activation maps shown as multi track diagrams of six cardiomyocyte specific gene loci (*Mybpc3, Myh7, Tnni1, Cox6a2, Myl3, Rcan1*) spanning 140 Kbp each. The TSS of each index gene is shown in red arrows. Tracks showing (from top to bottom): The genomic track with exons and introns in blue; Activation track showing index gene fold change activation following dCas9-VPR targeting to the genomic site vs. non-targeting gRNA control, as measured by RT-qPCR normalized to *Gapdh* in FIB (Scale of activation for the track is shown in square brackets, average fold activation from each gRNA site in black bars, red dots over bars indicate standard error, all black bars have p<0.05 vs. control, and sites with non-significant p>0.05 activation are shown as pink bars on the negative scale for visibility, n=3 for all sites); ATAC-seq and H3K27ac ChIP-seq tracks are shown for FIB in greens and CM in blue. HiC maps for each locus are shown above, with aqua colored lines indicating insulation locus boundaries. (**b**) Scatter plot showing change in target gene fold expression following specific gRNA targeting in all six genes shown in (a) as a function of the gRNA distance from the gene TSS in Kbp. Each dot represents the average of n=3 replicates, only dots were expression vs. control p<0.05 were included. The plot shows a non-linear decay in activation as a function of the distance.

Together these data show that recruitment of multiple strong activation domain using dCas9-VPR is capable of strong induction of gene expression in inactive genes. This induction is generally strongest when targeting sites near the TSS, and the degree of activation drops non-linearly as the distance from the TSS increases. Nevertheless, significant induction was still achieved from multiple non-proximal sites up to 70 Kbp away from the TSS.

### Genes can be activated across insulating loci from arbitrary genomics sites

Topologically Associating Domains (TADs) have been suggested to specify regulatory microenvironments for enhancer-promoter interaction (Dixon et al., 2012; Lupiáñez et al., 2015; McCord et al., 2020; Nora et al., 2012; Sexton and Cavalli, 2015). We therefore asked whether dCas9-VPR activation of genes behaves differently when crossing such insulating loci. To this end, we performed Hi-C to produce the first Rat fibroblast interaction maps and identified TAD insulating loci at 5Kbp resolution using an adaptation of the insulation score (Crane et al., 2015a) (Fig. 2a). Since the activation sites were chosen solely based on their distance from the TSS, most of the targeting sites fell within the same insulating locus as the index gene, however, several targeted sites fell beyond the gene-containing insulating locus. Although these sites tended to be far from the TSS, they were in some cases still successful in upregulating the target gene (Fig. 2a). We show for instance, that an activation site positioned +40 Kbp from the *Mybpc3* gene TSS, falling withing a different insulating locus than the gene promoter, increased this gene activation by 23±2.8-fold relative to non-targeting gRNA. To quantify the effect of insulation, without being confounded by the distance-activation relationship we looked for pairs of sites positioned at the same distance from the TSS of the index gene but lying either within or outside the insulating locus of the gene. We could find 11 such pairs, located in the *Mybpc3, Cox6a2*, and *Tnni1* loci, where the TSS is relatively close to the insulation boundary. The effect of insulation for activation from outside compared to activation from within the locus for pairs with similar distance from the TSS was modest and not statistically significant (8.75±0.89 vs 10.35±0.97-fold activation respectively, paired t-test p=0.28, Fig. S2). While the number of distance matched pairs we could find was relatively small, and we cannot exclude some insulation effect, we had 80% and 92% power to detect a 75% and 90% insulation effect respectively by insulating loci.

Next, we asked if any genomic site could serve as a CRE if strong activation domains were recruited to it. The gRNAs used for recruiting dCas9-VPR were chosen based on a pre-specified distance from the TSS to avoid any bias. Of the targeted genomic sites 127/149 (85.2%) induced significant activation (Table S1), showing that most genomic sites within 70 Kbp from the index gene could serve for activation. We used our mapping of open chromatin by ATAC-seq and H3K27 acetylation by ChIP-seq ((Golan-Lagziel et al., 2018), GSE102532) in FIB and in CM to examine the chromatin characteristics of the activation sites prior to dCas9-VPR recruitment. This analysis showed that 80% of gRNA sites that induced a significant change in gene expression targeted closed chromatin and mostly areas devoid of active enhancer and promoter histone acetylation mark H3K27ac in either FIB or CM (Fig. S1, Fig. S3). Since very few targeting sites were located in open chromatin, and since sites near transcription start sites, where activation is strong, tend to have open chromatin, we cannot reliably determine if targeting open chromatin would result in greater activation than targeting close chromatin, nor was it our aim. Nevertheless, our data clearly shows that recruitment of dCas9-VPR to non-regulatory sites lacking open chromatin or H3K27ac marks is sufficient to induce expression of distant genes. The absence of regulatory features at many of these sites even in CM, a cell type where these genes are highly expressed, further indicates that these sites do not function as endogenous regulatory elements. Together our results show that activation can occur by targeting naïve non-regulatory sequences and across insulating boundaries.

### Recruitment of activation domains result in epigenetic activation of the targeting site and the distant promoter

H3K27ac and H3K4me1 histone modifications were both shown to mark active enhancers and promoters (Creyghton et al., 2010; Local et al., 2018). Specifically, H3K27ac can differentiate between active and poised enhancers in mammalian cells (Creyghton et al., 2010). We tested whether recruitment of activation domains to genomic sites lacking such activation marks would result in histone-mark gain. To this end, two CM-specific genes, *Mybpc3* and *Cox6a2*, were targeted for activation in FIB from site located 20 Kbp downstream of their TSS. Both targeted distal sites have closed nucleosomes and lack significant H3K27ac active enhancer marking in both FIB and CM (Fig. 3a). Recruiting dCas9-VPR to these sites resulted in robust induction of *Mybpc3* and *Cox6a2* genes as measured by qRT-PCR (Fig. 3b). We then analyzed H3K4me1 and H3K27ac histone marks by ChIP-qPCR as a percentage of input at both the gRNA targeting site, 20 Kbp from the TSS, as well as at the gene promoter. Two primer pairs, ~300 bp apart, were used for each of these regions. This analysis showed that recruitment of a strong tandem activation domains to a distant non-regulatory site was sufficient to confer active regulatory element histone marks, H3K27ac and H3K4me1, at both the targeted site and at the promotor, compared to cells transfected with non-targeting gRNA (Fig. 3c).

**Fig. 3.**
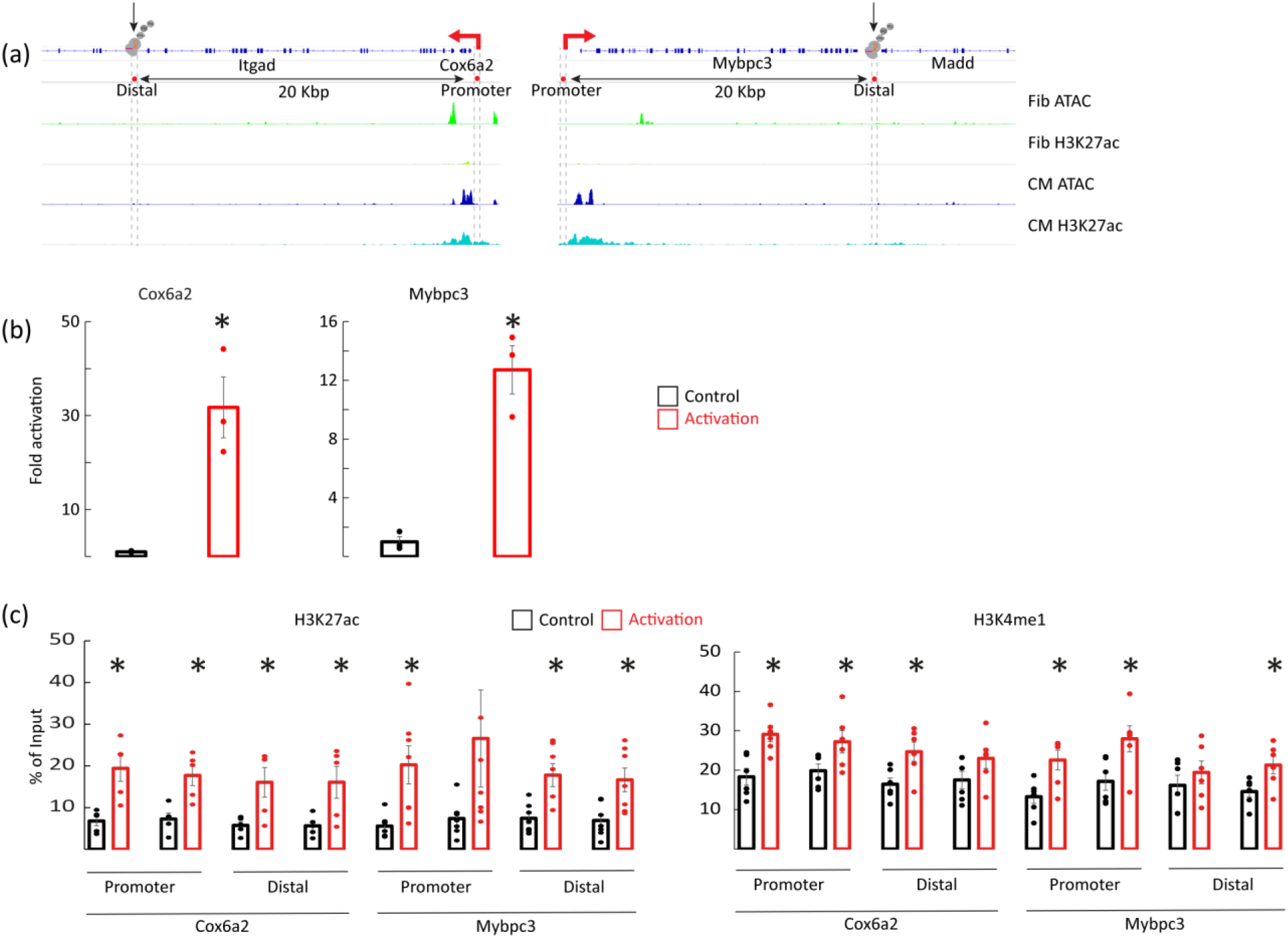
Activation from distal sites confers epigenetic marks at the activated site and at the gene promoter. (**a**) A diagram of the *Cox6a2* and *Mybpc3* loci with a genomic track showing exons and introns in blue and ATAC-seq and H3K27ac ChIP-seq track in CM and FIB. A distal locus, marked by a black arrow and cross-hatched lines, was activated by dCas9-VPR in FIB. We then assessed *Cox6a2* and *Mybpc3* activation and the epigenetic marks at the distal site of activation and at the promoter of these genes (red arrow). As shown, prior to activation the distal sites lacked ATAC and H3K27ac marks in both FIB and CM, and the promoter lacked such marks in FIB. (**b**) qRT-PCR results show that dCas9-VPR targeting the 20 Kbp distal site induces strong gene activation. Data is shown as fold activation vs. non targeting gRNA control normalized to *Gapdh* (n=3, * p<0.01). (**c**) ChIP-qPCR analysis of H3K27ac (left) and H3K4mm (right) histone marks shows increased chromatin modification of both the distal activation site and the gene proximal promoter following activation of the distal site. For each promoter and distal site two separate primer pairs were used, amplifying regions ~ 300 bp apart. (n=5-8 for each primer pair) in N=2-3 independent experiments (*p<0.05).

### Recruitment of endogenous transcription factor activation domains shares many properties with recruitment of viral activation domains

The activation domains VP64 and Rta in dCas9-VPR are derived from viral genes. We wanted to examine if the properties we observed with dCas9-VRP also apply to non-viral activation domains. The cardiac transcription factors *Gata4, Nkx2-5*, and *Tbx5* often co-occupy the same enhancers and their activation domains were previously identified (Morrisey et al., 1997; Ranganayakulu et al., 1998; Zaragoza et al., 2004). We used them to create dCas9-*Gata4-Nkx2-5-Tbx5* (dCas9-GNT) endogenously based CRISPRa tool (Fig. 4a). We selected 18 gRNA target sites covering a distance up to ± 30 Kbp from the TSS of the *Tnni1* and the *Cox6a2* genes, and mapped the activation induced by dCas9-GNT from these sites in FIB (Fig. 4b). This analysis shows that activation by dCas9-GNT behaved similarly to activation by dCas9-VPR in that dCas9-GNT could activate genes from a distance, even when targeting ‘naive’ genomic sites lacking open chromatin or active enhancer marks, and that the activation tended to be stronger when targeting sites near the promoter of the genes. Like dCas9-VPR the dCas9-GNT could activate these gene across an insulation boundary, and for example we could achieve a 6.4±1 and 2.88±0.23 fold activation of the *Tnni1* and *Cox6a2* genes respectively from sites located 30 Kbp upstream of these genes and in a different insulating locus. Finally, we chose a site located 15 Kbp upstream of the *Myh7* that lacked open chromatin or H3K27ac marks in FIB for activation by dCas9-GNT (Fig. 4c-d). The ChIP-qPCR showed that recruitment of dCas9-GNT to this site was sufficient to confer the active enhancer histone mark H3K27ac to both the targeted site and the *Myh7* promotor (Fig. 4e). Together these data show that endogenous transcription factor activation domains share the properties we observed with the viral derived activation domains in VPR.

**Fig. 4.**
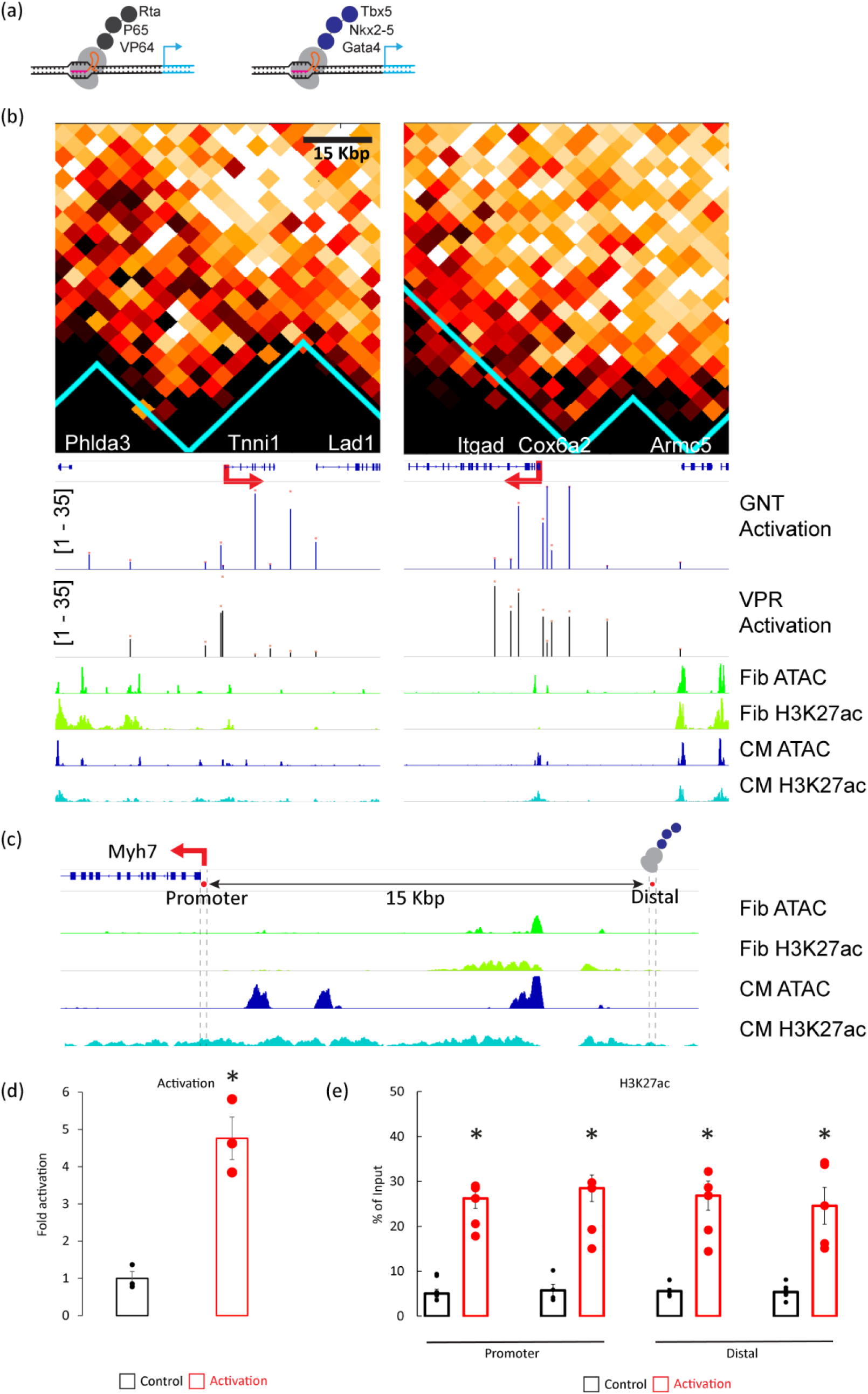
Activation with endogenous activation domains has similar features to activation with viral derived activation domains. (**a**) Diagram of CRISPR activators composed of dCas tethered to the activation domains of VP64, p65 and Rta (dCas9-VPR, left) or dCas tethered to the activation domains of the endogenous cardiac transcription factors Gata4, Nkx2-5, and Tbx5 (dCas9-GNT, right) (**b**) Activation maps as multi track diagrams of two cardiomyocyte specific gene loci (*Tnni1, Cox6a2*) are shown in a 70 Kbp window around the TSS. Tracks showing (from top to bottom): The genomic track with exons and introns in blue; Activation tracks showing target gene average fold activation following dCas9-GNT in blue bars or activation with dCas9-VPR in black bars targeted to the genomic site vs. non-targeting gRNA control, as measured by RT-qPCR normalized to *Gapdh* in FIB (Scale of activation for the track is shown in square brackets, red dots over bars indicate standard error, only bars with p<0.05 vs. control are shown, n=3). Tracks of ATAC-seq and H3K27ac ChIP-seq are shown for FIB in green and CM in blue. Diagram shows similar activation by dCas9-GNT and dCas9-VPR from distal sites lacking open chromatin or H3K27ac marks. (**c**) A diagram of the *Myh7* locus with a genomic track showing exons and introns in blue and ATAC-seq and H3K27ac ChIP-seq track in CM and FIB. A distal site, 15 Kbp from the TSS, marked by a red dot and cross-hatched lines, was activated by dCas9-GNT in FIB. We then assessed the H3K27ac marks at the distal site of activation and at the promoter of these genes (red arrow). (**d**) qRT-PCR results show that dCas9-GNT targeting the 15 Kbp distal site induces strong activation of *Myh7* in FIB. Data is shown as fold activation vs. non targeting gRNA control normalized to *Gapdh* (n=3, * p<0.005). (**e**) ChIP-qPCR analysis of H3K27ac activation marks shows increased chromatin modification of both the distal activation site and the gene proximal promoter following activation of the distal site by dCas9-GNT. For each promoter and distal site two separate primer pairs were used, amplifying regions ~ 300 bp apart. (n=5-8 for each primer pair in N=2-3 independent experiments, *p<0.05).

### Recruitment of repression domain downregulates gene expression in a distance-dependent manner from multiple genomic loci

Next, we examined whether repression has similar properties to activation. We used dCas9 fused to the endogenous repression domain KRAB and asked if the same targeting sites used in FIB for gene activation with dCas9-VPR could be used for repression with dCas9-KRAB in CM, where these genes are highly expressed. Specifically, the *Myh7* and *Mybpc3* loci were targeted in CM from sites up to 70 Kbp away from their TSS. We used some of the same gRNAs used in FIB for activation and elicited significant activation in FIB. In total, 17 targeted sites were studied for their effect on gene repression in CM (Table S2). The KRAB repression domain recruitment in CM showed a similar pattern to the one observed with VPR activation domain recruitment in FIB (Fig. 5a). A robust repression was elicited from sites near the target gene TSS, with a decline in repression when targeting from a distance. Gene repression was achieved even when targeting sites with nucleosomal chromatin and with no active enhancer marking (Fig. S4), and like in activation, repression of target genes was achievable by recruitment of the repression domain to genomic loci residing in different insulating locus. For example, targeting a site 70 Kbp upstream of the *Mybpc3* gene TSS induced robust downregulation of 53% compared to non-targeting gRNA (n=3, p<0.05). By plotting gene fold activation in FIB as a function of percentage of repression in CM, elicited by the same gRNAs, we show that activation and repression, using CRISPRa and CRISPRi respectively, are weakly correlated with Spearman’s Rho 0.34 (2 tailed p=0.176, n=17) (Fig. 5b).

**Fig. 5.**
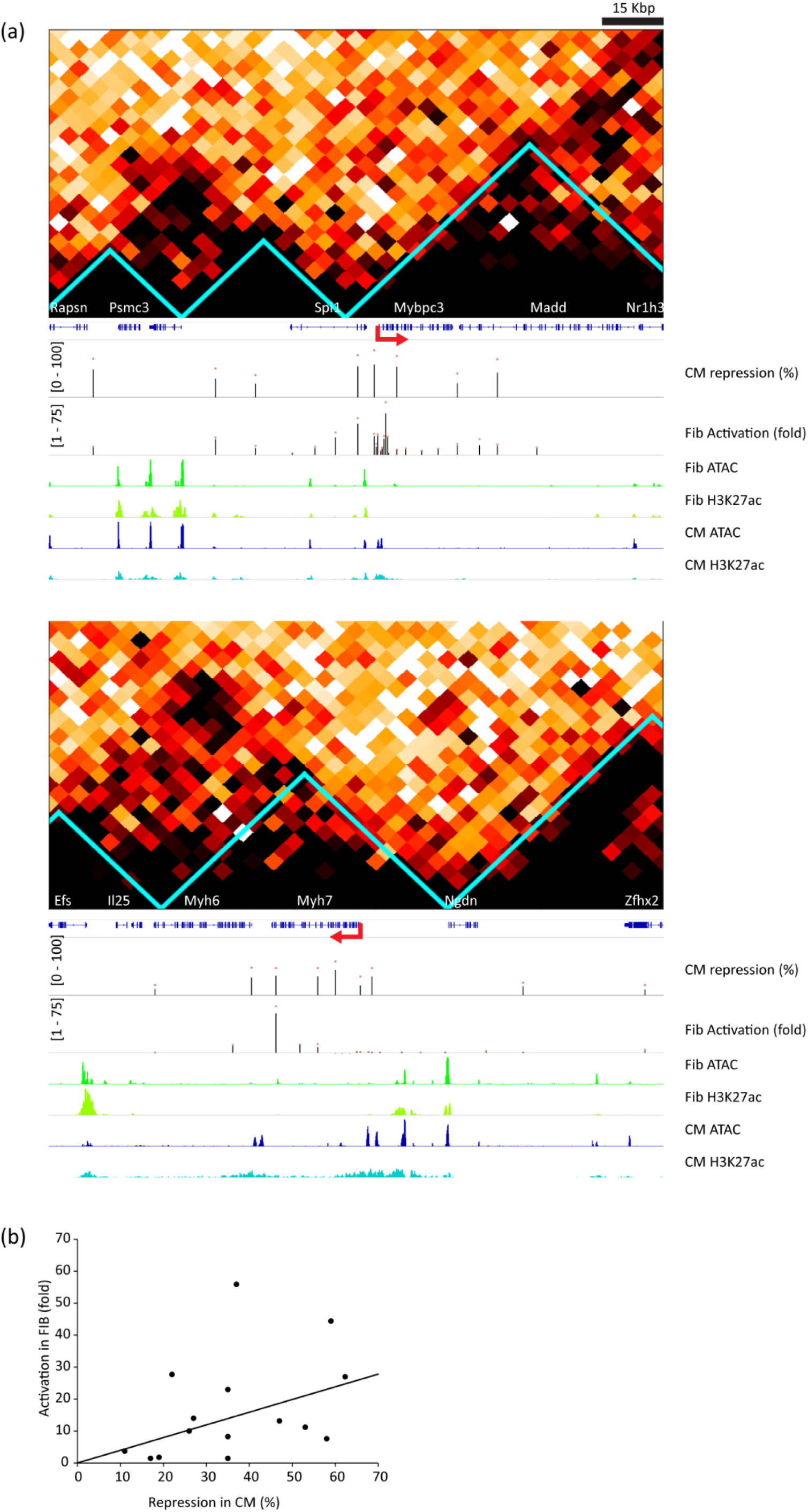
Recruitment of repression domain represses gene expression in a distance dependent manner from multiple genomic loci. (**a**) Multi track diagrams of two cardiomyocyte specific gene loci (*Mybpc3, Myh7*) spanning 140 Kbp each. TSS of each gene is shown in red arrows. HiC maps for each locus are shown, with aqua colored lines indicating insulation boundaries. Tracks showing (from top to bottom): The genomic track with exons and introns in blue; Repression track showing index gene % repression in CM following dCas9-KRAB targeting to the genomic site with targeting vs. non-targeting gRNA control, as measured by RT-qPCR normalized to *Gapdh* (Scale of repression 0-100% in square brackets, average % repression from each gRNA site in black bars, red dots over bars indicate standard error, only bars with p<0.05 vs. control are shown, n=3). Activation track showing the same index gene activation in FIB following dCas9-VPR targeting to the genomic site vs. non-targeting gRNA control, as measured by RT-qPCR normalized to *Gapdh* in (Scale of activation for the track is shown in square brackets, average fold activation from each gRNA site in black bars, red dots over bars indicate standard error, only bars with p<0.05 vs. control are shown, n=3). ATAC-seq and H3K27ac ChIP-seq tracks are shown for FIB in green and CM in blue. Diagram shows that like activation, the repression can be achieved at a distance, by targeting non regulatory chromatin, and can cross insulation boundaries. (**b**) Scatter plot of gene fold activation in FIB by dCas9-VPR as a function of % repression by dCas9-KRAB in CM as measured by RT-qPCR for multiple gRNA targeting sites in the *Mybpc3* and *Myh7* loci. Each dot represents the average of n=3 measurements in CM and FIB. Plot show activation and repression from these sites are generally correlated. A linear regression line is shown (Spearman’s Rho 0.34, n=17).

Next, we evaluated the repression of the *Myh6* gene, that encodes for cardiac α-myosin. We compared the degree of *Myh6* gene repression induced by targeting the promoter of *Myh6* or a distal site, 6 Kbp upstream of *Myh6* promoter, with dCas9-KRAB in CM (Fig. 6a). This upstream distal site resides inside the nearby *Myh7* gene. While the proximal promoter site has open chromatin in CM, the targeted distal site does not have this feature of a regulatory site, based on the ATAC-seq data (Fig. 6a). Gene expression analysis by qRT-PCR shows that targeting either the promoter or the distal sites resulted in significant gene repression (n=3, p<0.05) (Fig. 6b). In addition, we confirmed the repression by dCas9-KRAB using single molecule mRNA Fluorescent in-situ hybridization (smFISH). The quantification of smFISH signal in the nuclear transcription sites and of the cytoplasmic signals are surrogate measurements for the transcription rate and the mRNA levels respectively (Bahar Halpern et al., 2015; Lewis et al., 2018), and we have previously used this approach to study the transcription of the *Myh6* gene (Lewis et al., 2018). Representative images of the *Myh6* smFISH analysis in control CM and after repression from the distal site or from the promoter site show marked reduction in both nuclear sites and cytoplasmic signal (Fig 6c). Quantification of the smFISH images by FISH Quant tool (Mueller et al., 2013) showed significant reduction in both the cytoplasmic and transcription site smFISH signal by targeting either the distal or the promoter sites (Fig. 6d-e). These data confirm our qRT-qPCR analysis and show at the single cell level that recruitment of dCas9-KRAB to a distal site with closed chromatin can reduce the transcription rate of the targeted gene.

**Fig. 6.**
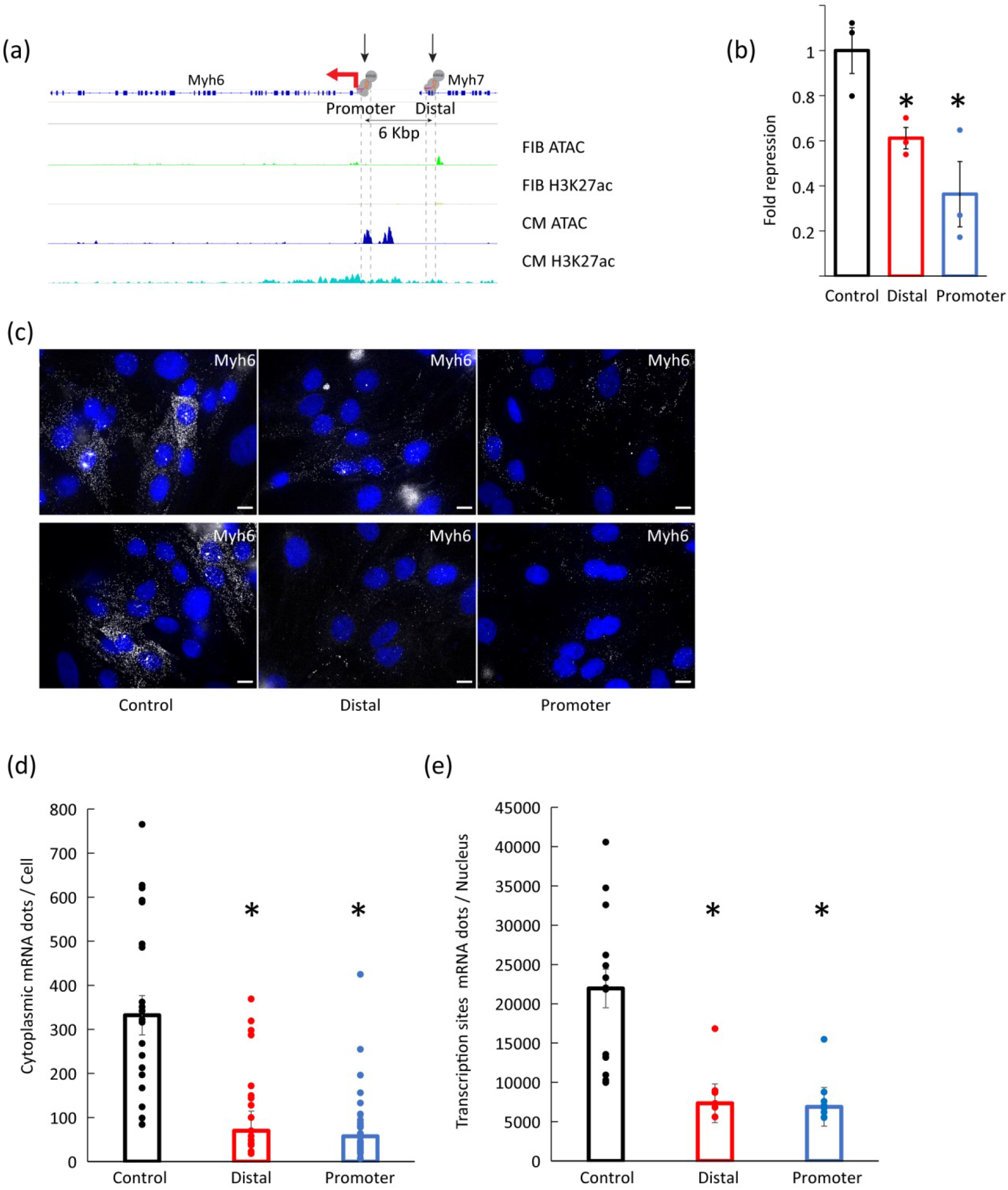
Repressor domain recruitment to distal non-functional genomic region in CM downregulates gene expression by inhibition of transcription. (**a**) Multi track diagram of *Myh6* gene locus, with the TSS of *Myh6* gene marked by red arrow. Tracks showing (from top to bottom): The genomic track with exons and introns in blue; ATAC-seq and H3K27ac ChIP-seq tracks are shown for FIB in greens and CM in blue. dCas9-KRAB was recruited by specific gRNAs to either the *Myh6* promoter or to a 6 KBp upstream distal site, marked by black arrows and cross-hatched lines. The distal site lacks ATAC open chromatin marks. (**b**) Assessment of *Myh6* gene repression following targeting of either the distal or promoter sites compared to non-targeting gRNA control by RT-qPCR normalized to *Gapdh* showing that recruitment of dCas9-KRAB to either site can significantly repress *Myh6* expression (bars show mean ±SE, n=3, *p<0.05). (**c**) Representative smFISH images of *Myh6* mRNA in CM show reduced cytoplasmic signal of mature mRNA, and reduced nuclear spots indicating transcription sites in CMs transduced with dCas9-KRAB recruited to either the distal or promoter regions (smFISH signal in white, Nuclei DAPI signal in blue, scale bar = 10 μm). (**d**) Quantification of cytoplasmic smFISH signal showing reduced mRNA levels when targeting either the distal or the promoter sites (n=20-30 cells, N=3 biologic replicates, *p<0.001) (**e**) Quantification of transcription sites smFISH signal indicating reduced mRNA transcription rate when targeting either the distal or the promoter sites (n=7-15 cells, N=3 biological replicates, *p<0.001).

## Discussion

We examined whether recruitment of transcription factor effector domains to arbitrary genomic sites is sufficient to establish a CRE at that site. We studied the epigenetic and transcriptional consequences of such recruitment as well as the effects of distance from a distal gene on the transcriptional output. We show that activation domain recruitment to naïve genomic sites, devoid of open chromatin or active enhancer chromatin marks, results in acquisition of such active histone marks both at the targeted site, the distant gene promoter, and in induction of distant gene expression. The distance between the enhancer and the cognate gene affects its strength non-linearly. Repression behaved in a similar manner to activation, and the targeting of non-regulatory regions could repress gene expression from a distance with a non-linear distance dependence and could cross chromatin contact insulation boundaries.

We tested two types of activator domain combinations - the VPR composed of the viral VP64 and Rta and of the p65 activation domains, and GNT-composed of *Gata4*, Mkx2-5, and *Tbx5* endogenous cardiac transcription factor activation domains, and both combinations could activate genes from a distant arbitrary genomic site. We also show that such activation domain recruitment to a distant site is sufficient to epigenetically mark both the distant site and the promoter with H3K27ac and H3K4me1. Likewise, the recruitment of the endogenous KRAB domain to an arbitrary site could repress a gene from a distance. The enhancer creation model suggests that certain transcription factors, termed pioneer factors, can bind to sequence specific motifs in nucleosomal chromatin to initiate enhancer assembly (Siegel and Sisler, 1963). However, the minimal required molecular components for enhancer assembly and series of events leading to enhancer activation are not fully understood (Field and Adelman, 2020). Our data supports the general conclusion that the recruitment of effector-activation or repressor domains to an arbitrary genomic site is sufficient to establish a CRE (enhancer of repressor) in the site and control the expression of genes from a distance. With ~1600 transcription factors encoded in the mammalian genome (Lambert et al., 2018), and the enormous number of potential factor combinations it remains to be studied which domain or domain combination are sufficient. Nevertheless, as many effector domains can interact with the same general co-factor, co-regulator, or histone modifying enzyme (Frietze and Farnham, 2011), it is likely that the ability to control the expression of genes from a distance is a general feature of many transcription factor effector domains.

We functionally show for multiple loci that genes can be activated from a distance, but the degree of activation declines nonlinearly with the distance of the activation sites from the gene. An inverse relationship between distance and the activity of an enhancer was also observed in an imaging-based study that showed that increasing the distance between the enhancer and promoter in a reporter construct in flies from 6.5 to 9 Kbp reduced the size and the timing of transcriptional bursting with more than 50% reduction in the level of the total output (Yokoshi et al., 2020). An emerging model proposes that regulatory elements may have both an enhancer and promoter functions, as enhancers and promoters share many molecular features, and gene promoters can act as enhancers for other genes (Andersson and Sandelin, 2020; Arnold et al., 2013). Our data supports this model and shows that an activation domain complex that serves as a strong promoter can exert a transcriptional control as it is moved further from the gene, albeit with a non-linear loss of activity. We observed an inverse distance - activation relationship in all the six loci studied, but there were inter- and intra-loci variability in the degree of activation and in the effect of distance. It is likely that additional layers of complexity beyond the chromatin accessibility, H3K27 acetylation, and chromatin three-dimensional contacts we examined in these loci contributes to the variability. Such factors include DNA methylation, additional histone modifications, non-coding RNAs, as well as additional recruitment of coactivators and co-repressors to sites in these loci. It is also likely that some of the variability is due to variation in the gRNA on-target efficiency in recruiting dCas9.

We show that activation can cross chromatin insulation boundaries. The role of TADs in mediating enhancer-promoter functional relationships is not entirely clear, and ablation of TAD structure can sometimes have minor effect on gene expression and enhancer activity for most genes (Akdemir et al., 2020; Despang et al., 2019; Ghavi-Helm et al., 2019; Rao et al., 2017; Schwarzer et al., 2017; Williamson et al., 2019). The results of our study are also in line with a high-resolution promoter interaction analysis that showed that about a third of significant promoter-putative regulatory element interactions occurred across TAD boundaries (Javierre et al., 2016). TADs and insulation events are stochastic such that any specific locus is insulated in only a subpopulation of cells, and transcription is stochastic as well (Bohrer and Larson, 2021). We used smFISH to examine the repression of *Myh6* in individual cells and saw that while many cells were affected, there was a significant variability in the degree of repression between individual cells (Fig. 6). It is therefore quite possible that most of the activation or repression effect we measure comes from the subpopulation cells in which little or no insulation was occurring.

Previous CRISPRa and CRISPRi studies were focused on the developed of the tools, showed that genes can be activated or repressed by targeting their promoter or their enhancers, and demonstrated that regulatory elements can be identified by tiling assays (Chavez et al., 2016; Cheng et al., 2013; Fulco et al., 2016; Gilbert et al., 2014, 2013; Kearns et al., 2014; Klann et al., 2017; Konermann et al., 2015; Lin et al., 2015; Maeder et al., 2013; Mali et al., 2013; Perez-Pinera et al., 2013; Simeonov et al., 2017; Tanenbaum et al., 2014; Xie et al., 2017). Here our aim was different, and we used CRISPRa and CRISPRi to identify the requirements and consequences of *de novo* CRE generation, and to identify the effects of the position of the CRE on the transcriptional output of distant genes. In addition to addressing these questions, our study has important implications for the use and interpretation of CRISPRa and CRISPRi experiments. In agreement with the previous studies, we found that for most loci the strongest activation and repression was achieved by targeting genes from sites near their TSS. Yet, activation at levels that are comparable to those achieved in previous studies that targeted gene promoters (Cheng et al., 2013; Li et al., 2020; Maeder et al., 2013; Mali et al., 2013; Perez-Pinera et al., 2013; Tanenbaum et al., 2014), could also be achieved at a distance. Therefore, one implication of our study is that multiple gRNA sites can be tried when aiming to activate or repress a gene, including sites that are not inside proximal promoters. We also show that the ability to activate or repress a gene from a distant site does not necessarily indicate that the targeted site is an endogenous regulator.

In summary, using an unbiased approach we show here that recruitment of effector domains to a single naïve genomic site can control the expression of a distant gene. When the distance between the enhancer and the promoter increases, the enhancer’s activity decreases. We speculate that cells overcome this limitation by combining multiple sites to form a larger and stronger enhancers (Hnisz et al., 2013), and by combining multiple enhancers to control each gene.

## Material and methods

### Cell Culture

Primary cultures of neonatal rat ventricular cardiomyocytes (CM) were isolated as previously described using the neonatal cardiomyocyte isolation system (Worthington Biochemical Corporation) from 1-3 day old Fischer rat pups (Golan-Lagziel et al., 2018). Cardiomyocyte fraction was purified by density centrifugation in Percoll (Sigma-Aldrich, St. Louis, MO) gradient. 3X10^6^ live cardiomyocytes were plated on 10cm dishes or 6 well plates, pre-coated with Cultrex Basement Membrane Extract (BME; Trevigen) diluted with serum-free DMEM. The culture medium was replaced 24h after plating with serum free medium and cultured for an additional 48 hrs. Fischer rat fibroblasts (FIB) were acquired from ATCC (Rat2, CRL-1764). All animal experiments were performed in compliance with relevant laws and institutional guidelines and approved by the local animal ethics committees of the Technion, Israel Institute of Technology.

### gRNA synthesis

The CRISPOR tool (Concordet and Haeussler, 2018) was used to choose appropriate and specific gRNA target sites closed to each mapping point. The single strand DNA oligonucleotide templates were acquired from Integrated DNA Technologies (IDT). Guide RNAs were in-vitro synthesized using Engen gRNA synthesis Kit (New England Biolabs) according to the manufacturer instruction with synthesis of the double stranded DNA template and transcription of RNA in a single reaction. Guide RNAs were then purified using Monarch RNA Cleanup Kit (New England Biolabs T2040L) per manufacturer instructions.

### Plasmids and transfections

Plasmids containing dCas9-VPR (SP-dCas9-VPR was a gift from George Church (Addgene plasmid # 63798; http://n2t.net/addgene:63798; RRID:Addgene_63798)) (Chavez et al., 2015), dCas9-KRAB domain (pLV hU6-sgRNA hUbC-dCas9-KRAB-T2a-Puro was a gift from Charles Gersbach (Addgene plasmid # 71236; http://n2t.net/addgene:71236; RRID:Addgene_71236)) (Thakore et al., 2015) were acquired from Addgene.

FIBs were transfected with dCas9-VPR construct 24 hours post platting on 6 well plates using Polyjet transfection reagent (Bioconsult SL100688) per manufacturer instructions. 48 hours post cell plating, sgRNAs were transfected using Lipofectaime RNAiMAX transfection reagent (Thermofisher) per manufacturer instructions.

### Adenovirus production

The dCas9-KRAB fragment was inserted into pEnt3c (Thermofisher) using HiFi DNA assembly (New England Biolabs), followed by Gateway LR Clonase (Thermofisher) reaction delivery into pAd-V5 Gateway Adenovirus destination vector (Thermofisher). Viral production and amplification in HEK293 cells as previously described (Golan-Lagziel et al., 2018).

*Gata4-Nkx2-5-Tbx5* (GNT) activation domains (Mus musculus GATA4 amino acids 2-75, NKX2-5 amino acid 18-129, and Homo sapiens TBX5 amino acid 339-379) and were acquired as a gene block from Integrated DNA Technologies (IDT). GNT fragment was inserted into dCas9 vector using NEBuilder HiFi DNA assembly (NEB-E2621L). The dCas9-GNT fragment was inserted into pEnt3c (Thermofisher) using HiFi DNA assembly (New England Biolabs), followed by Gateway LR Clonase (Thermofisher) reaction delivery into pAd-V5 Gateway Adenovirus destination vector (Thermofisher). Viral production and amplification in HEK293 cells as previously described (Golan-Lagziel et al., 2018).

CM were transduced with adenoviral vectors 24 hours post plating, followed by transfection of a single sgRNA 48 hrs post plating using Lipofectaime RNAiMAX transfection reagent (Thermofisher).

### RNA extraction, cDNA synthesis and qPCR

RNA was extracted using NucleoSpin RNA extraction kit (Macherey-Nagel) according to the manufacturer instructions. FIB and CM RNA was extracted following dCas9-VPR and gRNA transfection, 72 hours post cell plating. Reverse transcription to cDNA was performed using all-in-one RT Mastermix (abm). The qPCR was performed using the Bio-Rad CFX90 thermocycler and SYBRgreen Master Mix (Rhenium AB-4385612) using gene specific primers and expression was normalized to *Gapdh*. Technical duplicates were used for each reaction in addition to the biological replicates. For sgRNA transfection experiments, negative control samples were treated with non-targeting sgRNA.

Primers used:

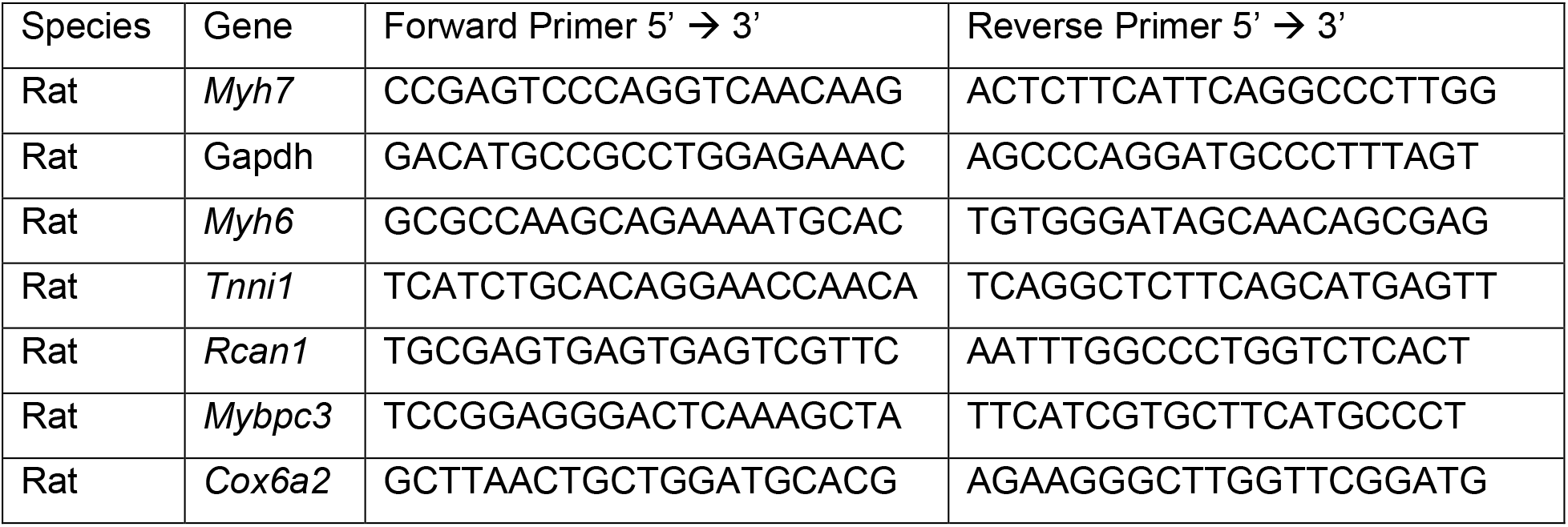

### RNA Single Molecule Fluorescence In-Situ Hybridization (smFISH)

Probe libraries were purchased from Biosearch Technologies as previously described (Lewis et al., 2018). Cells were washed with PBS, then fixed with a solution containing 3.5% formaldehyde and washed with a buffer containing 10% formamide (deionized; Sigma). Hybridization was performed in a buffer containing 10% formamide, 10% dextran sulphate, 2X SSC and desired probe at a concentration of 125 nM. Hybridization was done overnight in a humidified chamber at 37°C. Following hybridization, cells were washed twice with 10% formamide wash buffer for 30 min. Second wash was supplemented with 4’,6-diamidino-2-phenylindole dihydrochloride (DAPI) at a concentration of 1μg/ml for nuclear staining. Cells were then washed once with 2X SSC for 5 minutes and mounted on glass slides with 12 μl of Fluoromount-G (ThermoFisher Scientific), or with glucose oxidase – catalase anti-fade solution in case of Alexa-647 or Quasar-670 probe libraries. Slides were imaged with Axio Observer inverted fluorescent microscope (Zeiss) using an X-cite metal-halide light source and a high-resolution camera (Hamamtsu Orca R2), with an X63/1.4NA objective (Olympus). Exposure times for smFISH signal was between 300-600ms. Images were captured as a full thickness z-stack with a 0.24-0.30 μm section size. For qualitative analysis resulting images were imported into ImageJ, a Laplacian of Gaussian filter was applied to the smFISH channel using the LoG3D plugin, and a max intensity merge of the z-stack was acquired. Quantification of smFISH was performed using the FISH-quant MATLAB toolbox (Mueller et al., 2013).

### Chip-qPCR

Chromatin immunoprecipitation (ChIP) was performed by using MAGnify Chromatin Immunoprecipitation System (Invitrogen) with anti-Histone 3 acetyl K27 (H3K27ac) antibody (Abcam ab4729) and anti-Histone 3 k4 monomethyl (H3k4mm) (Abcam ab8895). Input controls were non-immune precipitated samples. ChIP-qPCR was done using a Bio-Rad CFX96 thermocycler. Data were calculated as a fraction of input chromatin.

### Hi-C

Hi-C was performed as described previously (Belaghzal et al., 2017). Briefly, ~10^6^ cells were cross-linked with formaldehyde, permeabilized and digested with DpnII. Next, sticky ends were filled with nucleotides including biotinylated dATP, followed by blunt-end ligation, cross-link reversal and DNA purification. This was followed by biotin removal from unligated ends, sonication, and pulldown of biotinylated fragments with streptavidin beads. Finally, the library was amplified, size selected and sequenced using 75bp paired-end sequencing on a NextSeq500.

The resulting 525,411,183 paired-end reads were processed as described previously (Lajoie et al., 2015). Briefly, read ends were independently iteratively mapped to DpnII restriction fragments based on the rat rn6 genome using bowtie2 (Langmead and Salzberg, 2012). The mapped reads were then filtered for artifacts and duplications, finally resulting in 285,170,763 valid unique read pairs. Valid read pairs were then binned into matrices, and the interaction matrices were filtered and balanced using Cooler (Abdennur and Mirny, 2020).

Insulation score was calculated following an approach previously described (Crane et al., 2015b), with slight modifications. Using 5kb bin resolution, the insulation score of position x was calculated as the mean of the square of all interaction frequencies between loci 100kb downstream with 100kb upstream of x. Discrete TAD boundaries were called using Scipy (Virtanen et al., 2020).

scipy.signal.find_peaks(insulation_score,prominence=0.00001,width=2).

### Statistics and informatic analysis

Most analysis steps are outlined in the text. Student’s two tailed t-test was used to compare the means of two groups unless otherwise specified. Analysis was performed using R (R Core Team (2016). R: A language and environment for statistical computing. R Foundation for Statistical Computing, Vienna, Austria. URL https://www.R-project.org/. HOMER annotatePeaks.pl was used for heatmap generation (Heinz et al., 2010). Estimated p values in the figures are result of double sided, unpaired t-student test, unless otherwise stated.

## Acronyms

CREs: *Cis*-acting regulatory elements
CRISPR: Clustered regulatory interspaced, short palindromic repeats
CAS9: CRISPR-associated protein 9
dCas9: nuclease-dead mutant of Cas9
CRISPRa: CRISPR activation
CRISPRi: CRISPR inhibition
TSS: transcription start site
KRAB: Kruppel associated box
ChlP-qPCR: chromatin immunoprecipitation followed by quantitative polymerase chain reaction
ATAC: assay for transposase-accessible chromatin
H3K27ac: histone H3 acetylated at lysine 27
H3K4me1: histone H3, monomethylated at lysine 4
FIB: fibroblasts
CM: cardiomyocytes
TAD: topologically associating domain
smFISH: single molecule fluorescence in-situ hybridization

## Data Availability

GSE102532

## Funding

IK was supported by the Israel Science Foundation (grant # 1385/20). NK was supported by the Azrieli Faculty Fellows program and Israel Science Foundation (grant# 1479/18).

## Conflict of Interest statement

The authors declare no conflicts of interest.

---

**Fig. S1.**
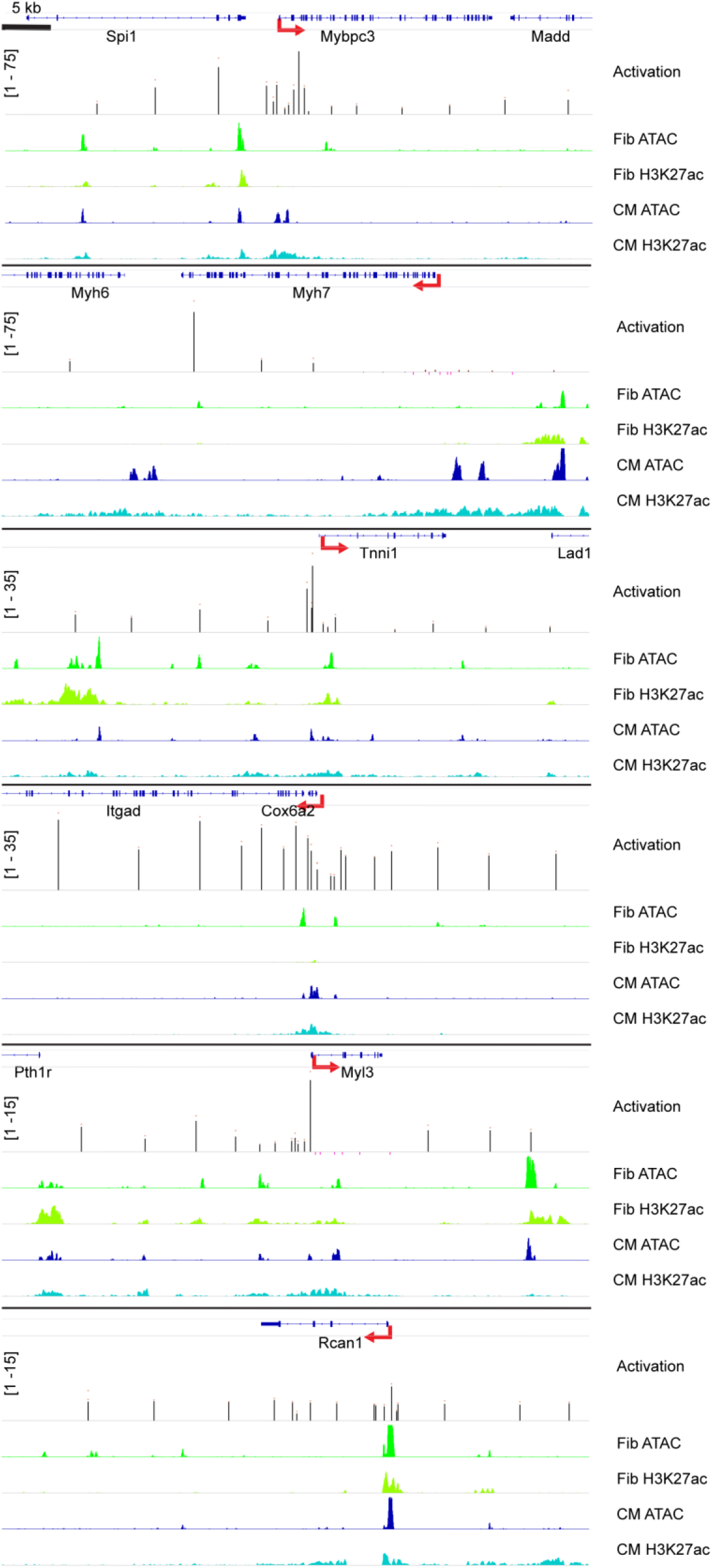
High resolution activation maps showing dCas9-VPR can activate genes from multiple sties. Activation maps shown as multi track diagrams of six cardiomyocyte specific gene loci (*Mybpc3, Myh7, Tnni1, Cox6a2, Myl3, Rcan1*) are shown at high resolution in a 50 Kbp window around the TSS. Tracks showing (from top to bottom): The genomic track with exons and introns in blue; Activation track showing index gene average fold activation following dCas9-VPR targeting to the genomic site vs. non-targeting gRNA control, as measured by RT-qPCR normalized to *Gapdh* in FIB (Scale of activation for the track is shown in square brackets, average fold activation from each gRNA site in black bars, red dots over bars indicate standard error, all black bars have p<0.05 vs. control, and sites with non-significant p>0.05 activation are shown as pink bars on the negative scale for visibility, n=3 for all sites); ATAC-seq and H3K27ac ChIP-seq tracks are shown for FIB in green and CM in blue.

**Fig. S2.**
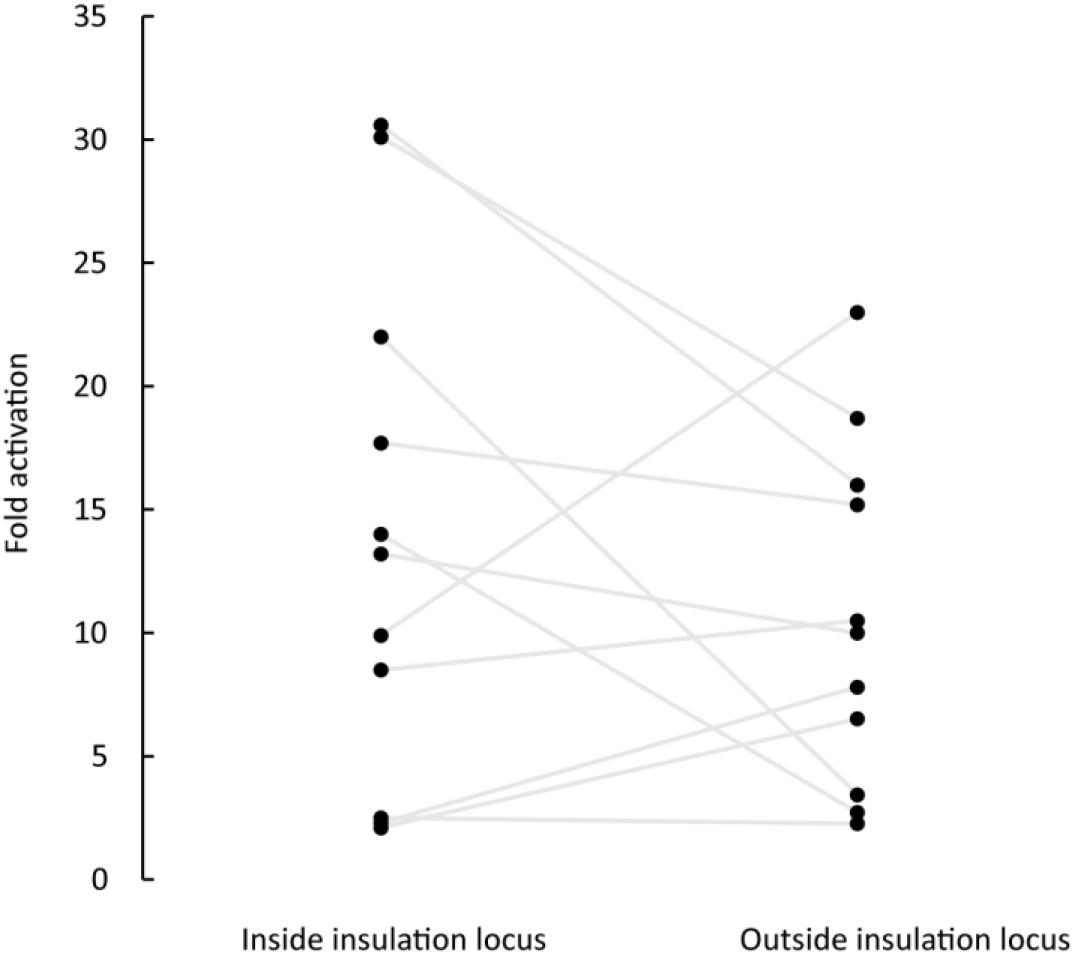
Recruitment of dCas9-VPR to sites outside insulation boundaries can induce the expression of genes. Comparison of the activation achieved from pairs of sites positioned at the same distance from the TSS of the index gene but lying within or outside the insulation locus of the gene showing no significant difference. (n= 11 pairs in the Mybpc3, Cox6a2, and Tnni1 loci, paired t-test p=0.28).

**Fig. S3.**
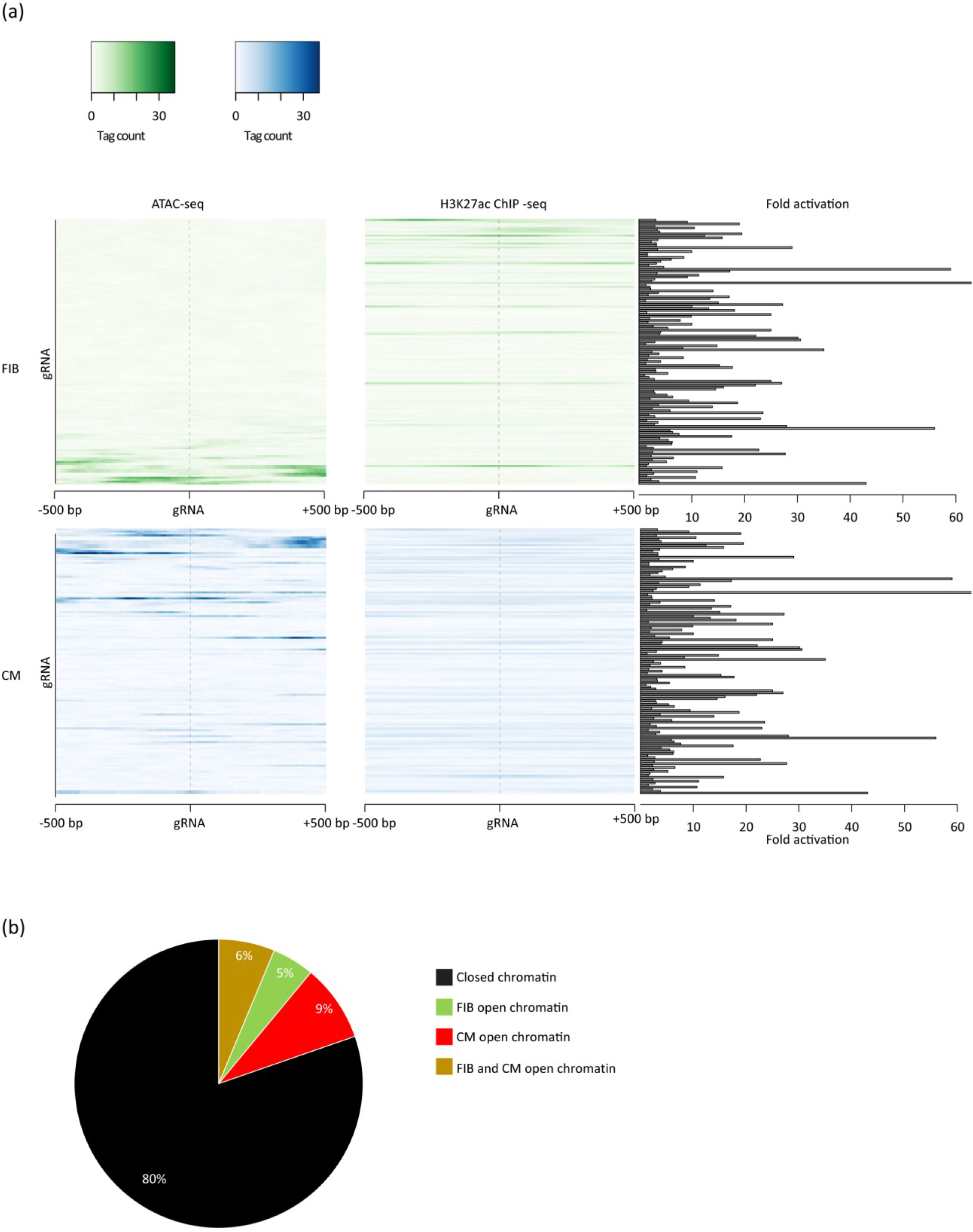
Recruitment of dCas9-VPR to non-regulatory sites lacking open chromatin or H3K27ac marks is sufficient to induce the expression of genes. (**a**) Heatmaps of ATAC-seq and H3K27ac signal in fibroblasts (FIB, green) and cardiomyocytes (CM, blue) centered on the gRNA targeting site and spanning ± 500 bp in each row. Bar plot on the right displays the fold-activation achieved by targeting the site in FIB, showing targeting sites lacking open chromatin or H3K27ac marks can result in strong activation. (**b**) Pie-chart showing the percent of gRNA sites falling on closed chromatin in both CM and FIB (black), chromatin that is open only in FIB (green), only in CM (red), or open in both FIB and CM (gold).

**Fig. S4.**
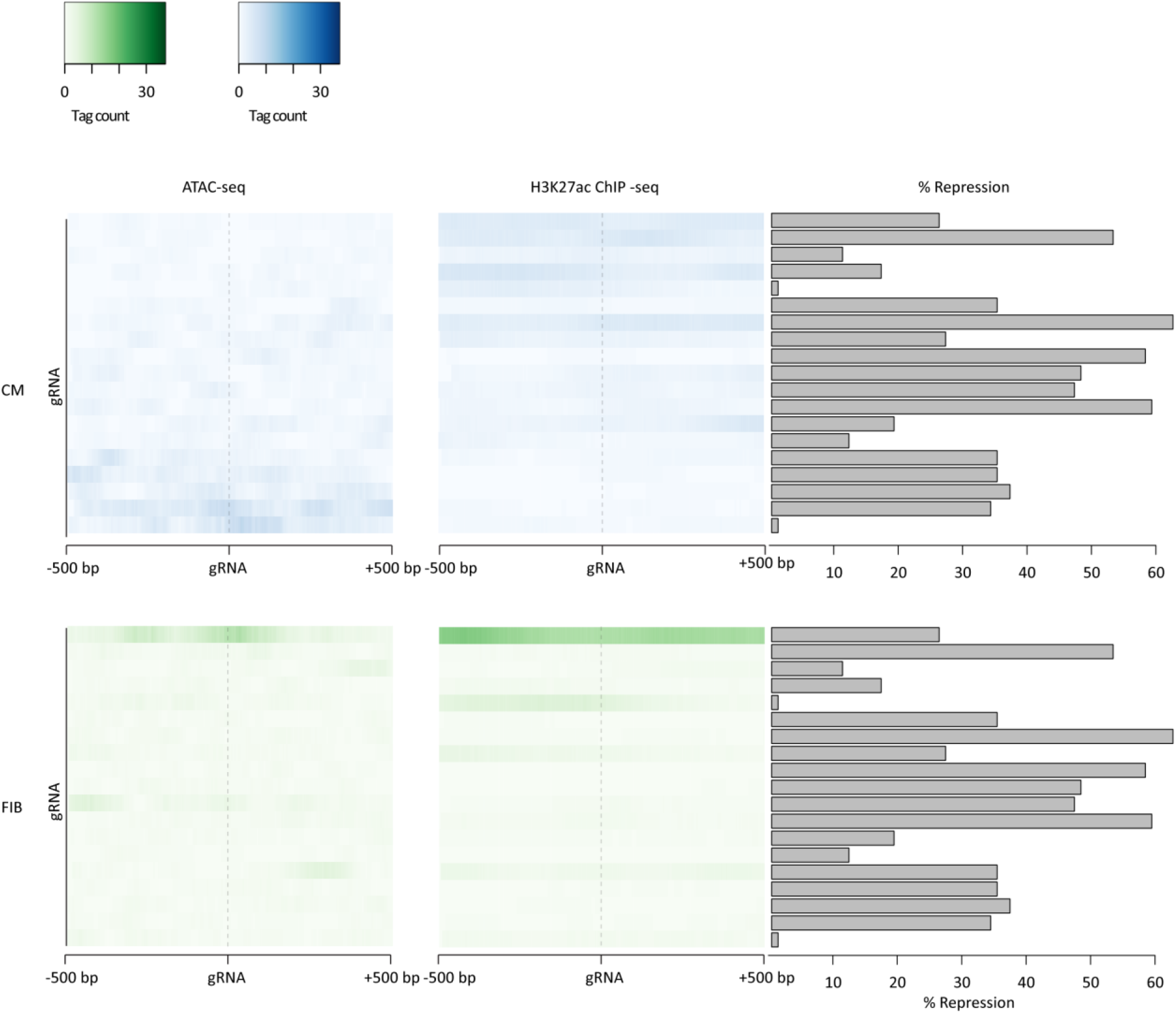
Recruitment of dCas9-KRAB to non-regulatory sites lacking open chromatin or H3K27ac marks is sufficient to repress the expression of genes. Heatmaps of ATAC-seq and H3K27ac signal in cardiomyocytes (CM, blue) and fibroblasts (FIB, green) centered on the gRNA targeting site and spanning ± 500 bp in each row. Bar plot on the right indicates the %-repression achieved by targeting the site in CM, showing that targeting sites lacking open chromatin or H3K27ac marks can result in strong repression.

## Notes

### Competing Interest Statement

The authors have declared no competing interest.

