## Supplemental table 1 for "Recruitment of transcriptional effectors by Cas9 creates cis regulatory elements and demonstrates distance-dependent transcriptional regulation"

Table S1

Table S1. Targeted sites as a function of distance from the TSS. Six CM-specific genes were targeted by dCas9-VPR activation complex in fibroblast. Check mark represents successful targeting with subsequent significant change in gene expression, Cross mark represents unsuccessful targeting, with no statistically significant change in gene expression.

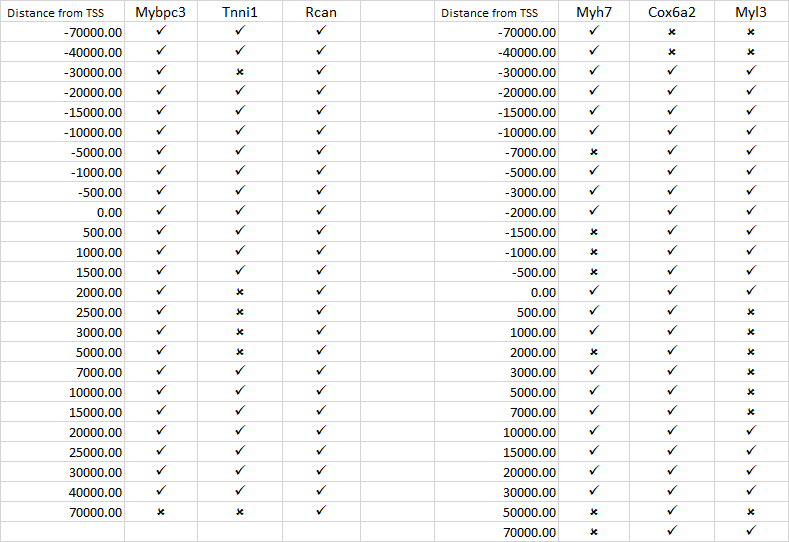
