## Supplemental table 2 for "Recruitment of transcriptional effectors by Cas9 creates cis regulatory elements and demonstrates distance-dependent transcriptional regulation"

Table S2

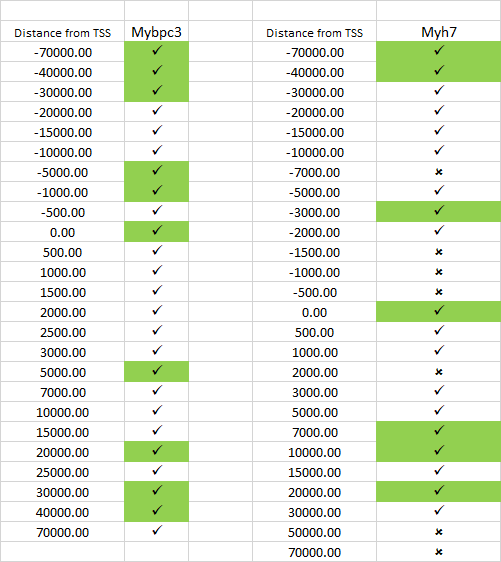

Targeting sites in the Mybpc3 and Myh7 loci with green labeled boxes representing genomic sites chosen for repression using dCas9-KRAB in CM.
